# An equivalence of prokaryotic pore forming proteins of Plasmodium triggers cellular dysfunction responsible for malaria pathogenesis

**DOI:** 10.1101/2020.08.01.230623

**Authors:** Abhishek Shivappagowdar, Swati Garg, Akriti Srivastava, Rahul S. Hada, Soumya Pati, Lalit Garg, Shailja Singh

**Affiliations:** Department of Life Science, School of Natural Sciences, Shiv Nadar University, Chithera, Gautam Buddh Nagar, UP, India; Special Centre for Molecular Medicine, Jawaharlal Nehru University, New Delhi, India; Gene Regulation Laboratory, National Institute of Immunology, New Delhi 110067, India

**Keywords:** Malaria, *Plasmodium falciparum*, Perforin like Proteins, necrosis, Blebbing, Calcium, HMGB1

## Abstract

Severe malaria caused by *Plasmodium falciparum* poses a major global health problem with high morbidity and mortality. The *P. falciparum* harbours a family of pore forming proteins (PFPs), known as perforin like proteins (PLPs), which are structurally equivalent to prokaryotic PFPs. These PLPs are secreted from the parasites and by interacting to host cells they contribute to disease pathogenesis. The severe malaria pathogenesis is associated with dysfunction of various barrier cells including endothelial cells. A number of factors, including PLPs, secreted by parasite contribute to the host cell dysfunction. Here in, we tested the hypothesis that the PLPs mediate dysfunction of barrier cells and might have a role in disease pathogenesis. We analysed various dysfunction in barrier cells following rPLP2 exposure and demonstrate that it causes an increase in intracellular Ca^2+^ levels. Additionally, rPLP2 exposed barrier cells displayed features of cell death including Annexin/PI positivity, depolarized mitochondrial membrane potential and ROS generation. We further performed the time lapse video microscopy of barrier cells and found the treatment of rPLP2 triggers their membrane blebbing. The cytoplasmic localization of HMGB1, a marker of necrosis, further confirmed the necrotic type of cell death. This study highlights the role of parasite factor PLP in endothelial dysfunction and provides a rational for the design of adjunct therapies against severe malaria.

## 1. Introduction

Malaria is probably one of the oldest life-threatening diseases caused by the protozoan parasites of the genus *Plasmodium*. The mortality due to malaria is extremely high, with approximately 85% of deaths that occur globally are among the children under the age of five [1]. While uncomplicated *Plasmodium falciparum* infection is highly treatable, the most severe complication, cerebral malaria, presents a significant problem for patients. Human cerebral malaria (HCM) is characterized by disruption of the blood-brain barrier, coma, seizures, and death. Patients that survive HCM are prone to persisting cognitive and neurological deficits after they have recovered from the infection. Although antimalarial drugs have shown efficacy in killing parasites, treatments to improve the outcome of HCM are lacking. Furthermore, the cellular mechanisms regulating HCM disease progression remain poorly understood. The difficulties associated with studying human patients during infection have limited our understanding of HCM and have created a need for additional preclinical research.

The pathophysiology of severe malaria is extremely complex and a field of intense research. The main mechanisms is host cell dysfunction due to sequestration of infected erythrocytes (IE), which bind to the vascular endothelium and induce activation of endothelial cells; activation of the immunological and systemic inflammatory responses; and coagulation dysfunction [2–4]. None of these mechanisms alone fully accounts for the pathogenesis of human cerebral malaria, and dynamic interactions occur among the three mechanisms, explaining the complexity of this potentially fatal infection [5]. The malarial parasite produces ligands, notably *Plasmodium falciparum* erythrocyte membrane protein-1 (PfEMP-1) which protrudes through the pRBC surface and binds to the receptors CD36, ICAM-1 (cerebral malaria), CS-A (placental malaria). This binding causes the sequestration (cytoadhesion) to the vascular endothelial cells, which is further increased upon agglutination with other non-pRBCs, resetting host leukocytes and platelets, finally slowing down the cerebral circulation and aggravating hypoxia through the coma. However, in the case of adhesion independent pathway, ligand-receptor specific interactions are not required. These pathways could activate secondary signaling cascade which includes induction of toxic local microenvironment, metabolic competition, the release of parasite products which act as toxins and other mediators [6]. All these factors cause cellular damage which could influence the expression of junctional proteins and also damage cells including microglia, astrocytes, neurons and other parenchymal cells [7].

Evidences have shown that the adhesion of pRBCs to the ECs can induce cell apoptosis via nitric oxide and oxidative stress [8]. Moreover, in the *in vivo* studies, the cytotoxic effector CD8+T lymphocytes are implicated in the development of murine CM by direct cytotoxicity against the ECs [9]. Recent studies by Tripathi et al [3] show that the trypsin-resistant membrane components of *Plasmodium falciparum*-infected red blood cells (Pf-IRBCs) and its soluble factors, contribute to the disruption of BBB integrity by leading to a decrease in HBMEC resistance. However, the exact soluble factors causing the disruption was not identified. Experimental cerebral malaria using *Plasmodium berghei* ANKA infection of C57BL/6 mice provides a setting for determining mechanistic details that is not feasible with human studies. In both humans and mice, parasitic infection initiates BBB disruption leading to fatal neurological dysfunction. The mechanism of disease onset involves a complex network of known effector cells and proteins. Studies have demonstrated that an influx of CD8+ T cells, macrophages, and neutrophils into the brain occurs during ECM [10]. In addition to immune infiltration, nitric oxide availability appears to play a role during ECM [8].

*Plasmodium falciparum* pore forming proteins (PFPs), known as perforin like proteins (PfPLPs), are the critical drivers of the parasite life cycle. They share structural similarity with bacterial PFPs both containing a central pore-forming domain comprising a bent and twisted β-sheet that is flanked by three clusters, usually comprising α-helices. Studies have shown that the human perforin secreted by CD8+ T cells cause BBB breakdown causing fatal edema [10]. In our previous work [11], we demonstrated that the highly conserved region of Plasmodium PLPs, MACPF domain can form pores on human erythrocytes leading to their lysis. Furthermore, they also induced an increase in intracellular calcium leading to premature erythrocyte senescence. Since it is known that the soluble factors of *P. falciparum* play an important role in the pathogenesis of CM, we hypothesized that the release of PLPs at the cerebral vasculature can lead to severe neurological dysfunction. Hence, we designed a model system using an epithelial as well as endothelial cells to study the pathogenesis of PLPs in the context of CM.

Our data indicate that recombinant PLP2 (rPLP2) induces an increase in intracellular Ca^2+^ levels in primary vascular cells, HUVEC and MDCK cells. Additionally, rPLP2 exposed ECs also displayed features of apoptotic cell death including exposure of phosphatidyl serine and propidium iodide positivity, loss of mitochondrial membrane potential, increased ROS generation and HMGB1 release. In conclusion, this study indicates the role of PLPs in CM, thus providing new insights into severe malaria pathogenesis.

## 2. Results

### 2.1. rPLP2 induces death of primary endothelial cells *in vitro*

It is well known that CDCs as well as MACPFs share an evolutionarily conserved structural fold as well as the mechanism of pore formation. The MACPFs, have a signature motif (Y/W)-X6-(F/Y)GTH(F/Y)-X6-GG and two transmembrane helices (CH1 and CH2). Although the MACPF domain is found mainly in eukaryotes, certain bacteria including Plu-MACPF from *Photorhabdus luminescens* also harbour these domains [12]. Our *in silico* analysis further indicates significant conservation of the MACPF domain between Plu-MACPF and PLP2 with two transmembrane helices present on either side of the β-sheet (Figure 1A).

**Figure 1:**
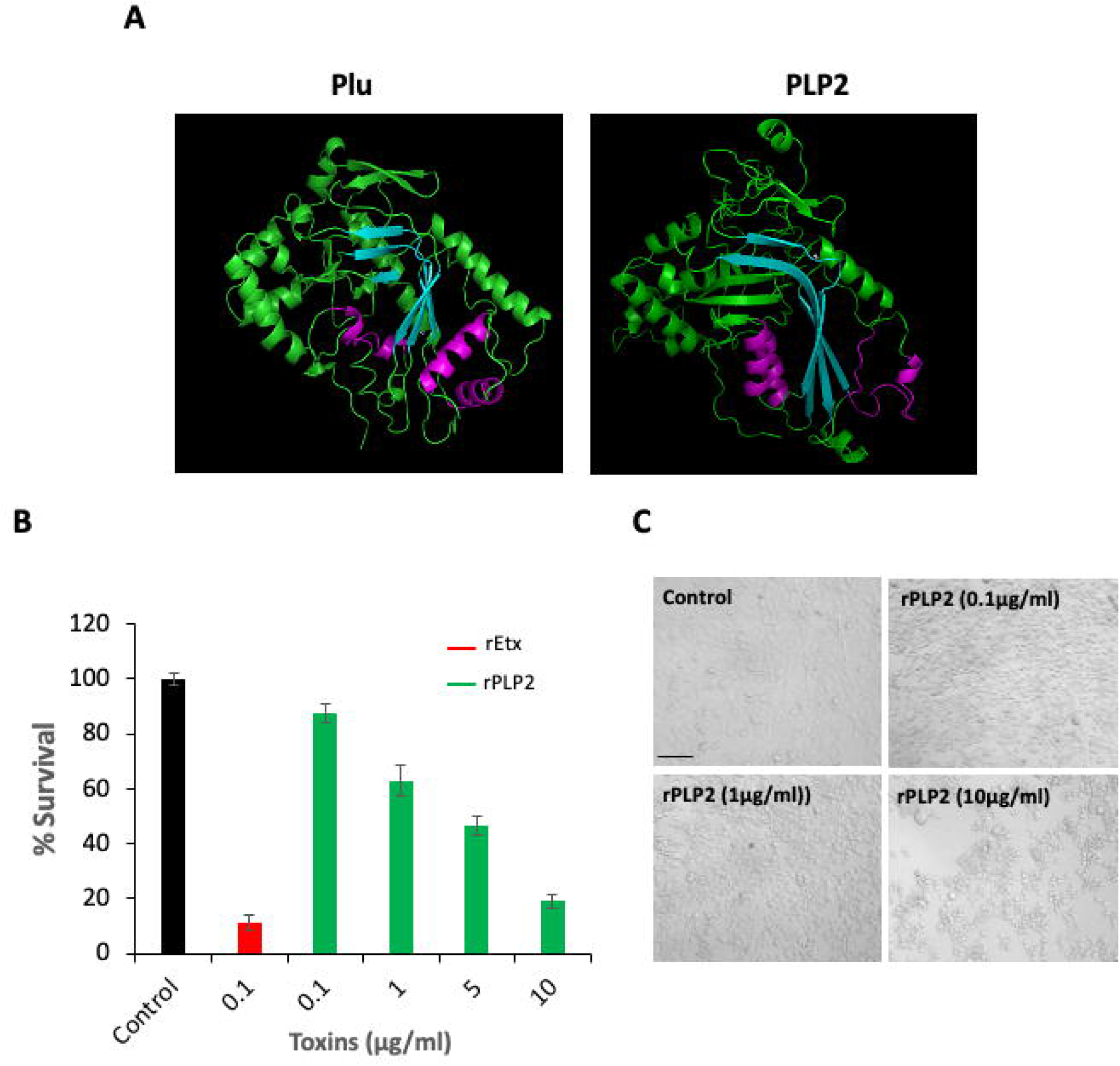
Activity of rPLP2. A) The tertiary structure conservation analysis between the MACPF domains of *Photorhabdus luminescens* Plu and *Plasmodium falciparum* PLP2, show high degree of similarity between them. The conserved regions are shown in the same colour. The central β-sheet is represented in cyan and the transmembrane hairpins (TMH) on the either side are shown in purple. B) MDCK cells were exposed with the indicated concentration of rPLP2 and the survival was affirmed by MTT assay. Error bar represents mean±SD. C) The microscopic analysis of the effect of rPLP2 on MDCK cells was monitored by incubating the cells with the different concentrations of the protein. Control cells were incubated with media only. Scale bar represents 20 μm.

To test the cytotoxic activity of rPLP2 on MDCK cells, we performed MTT assay *in vitro*. Briefly, rPLP2 was incubated with MDCK cells at different concentrations. The viability of the cells decreased in a dose-dependent manner with an increase in rPLP2 concentration. The lethal effect was observed starting from 100 ng/ml rPLP2 and the IC_50_ was found to be at approximately 5 μg/ml. The cells untreated with the protein were considered 100% survival (Figure 1B). A similar PFP of *Clostridium spp*, epsilon toxin (Etx) was used as control. To check for the morphological changes in MDCK cells, we performed light microscopy of rPLP2 treated cells. Similar to other toxins, the rPLP2 treated cells also displayed changes in the cellular architecture demonstrating membrane blebbing and vesiculation in a dose-dependent manner (Figure 1C). We next investigated the effect of rPLP2 on the shape and viability of MDCK and HUVEC cells during its toxic action. As shown (Figure 2A, 2B and Supplementary movie 1, 2) rPLP2 effect on the cells could be detected within few minutes of addition. Most of the cells developed swelling blebs that increased and grew over time. Swelling blebs are quasi-spherical protrusions typically formed on the cell surface after intense membrane injury. Although the role of blebbing under pathological conditions is still controversial, it has been proposed that it could represent a reservoir for the increased cellular volume of water and have a defensive function against cell lysis.

**Figure 2:**
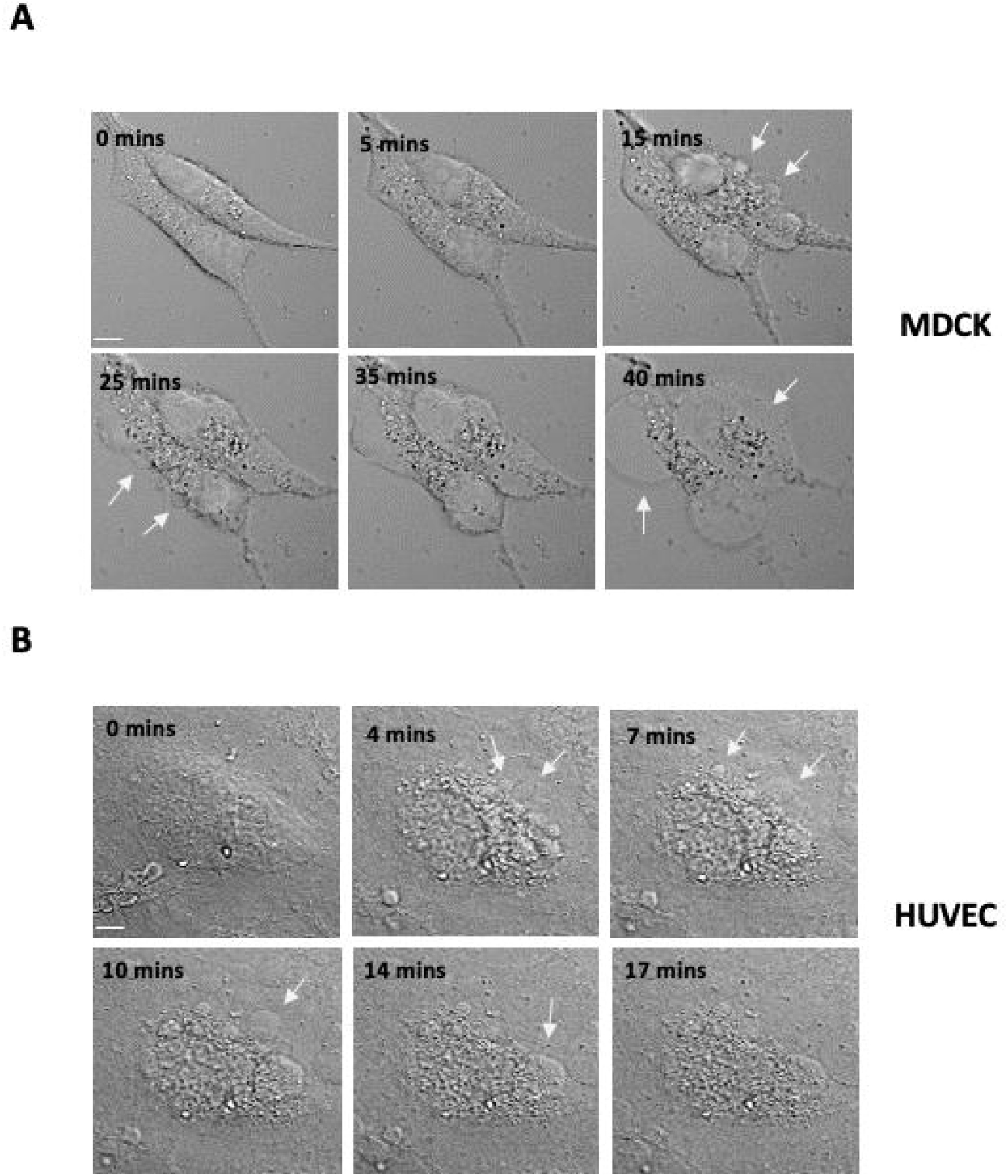
Membrane blebbing of the cells in response to rPLP2. The MDCK cells (A) and HUVEC (B) were treated with rPLP2 and the changes in the membrane architecture were observed in real-time by confocal microscopy. Selected images of DIC with time elapsed between frames in minutes (mins) are shown. Arrows indicate the blebbing of the cell membrane. Scale bar represents 2 μm.

### 2.2. rPLP2 dependent cell death involves an increase in the intracellular calcium levels

Influx of calcium is the primary consequence of impaired membrane, produce changes in cells that they can respond to by triggering the repairing mechanisms. The increase in Ca^2+^ levels can initiate cytoskeletal degradation, activate hydrolytic enzymes, impair energy production ultimately leading to cell death [13]. Calcium can enter the cytosol of endothelial cells through rPLP2 pores. Hence, we monitored rPLP2 mediated influx of calcium in primary endothelial cells. The rPLP2 treated MDCK and HUVEC cells displayed a clear increase in the intracellular Ca^2+^ levels compared to the control cells (Figure 3A and 3B). This increase could be due to the influx of extracellular calcium that occurs upon pore formation. To monitor more closely, we performed time lapse microscopy of HUVEC cells incubated with rPLP2. We observed a sudden increase in intracellular calcium of HUVEC cells following addition of rPLP2. The calcium influx further led to morphological changes in the cells leading to cell death (Figure 3C and Supplementary movie 3).

**Figure 3:**
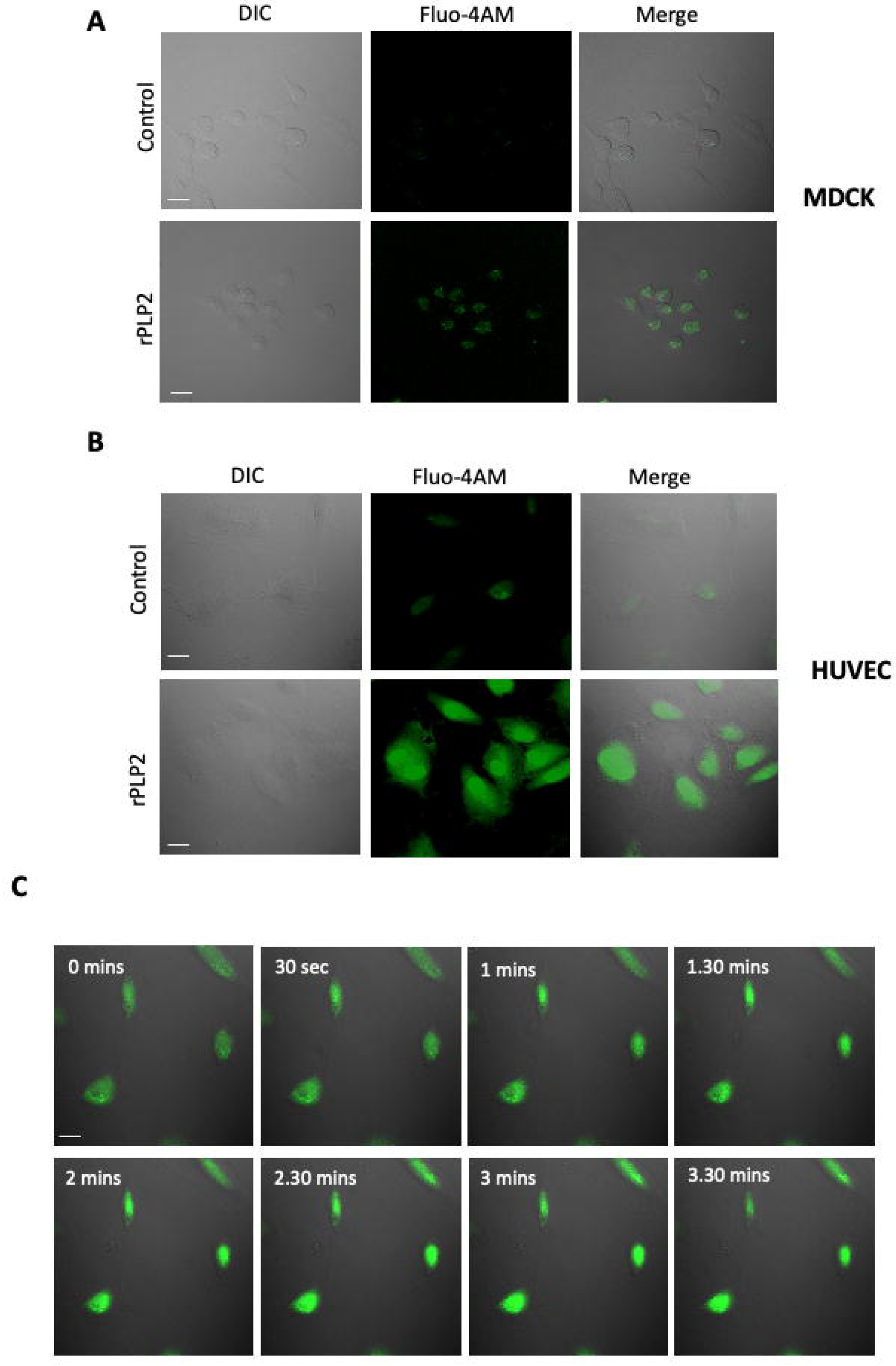
rPLP2 mediates calcium influx. Intracellular Ca^2+^ levels were monitored in MDCK (A) and HUVEC (B) cells loaded with Fluo-4AM in the presence or absence of rPLP2. Scale bars indicate 2 μm. C) Time-lapse video microscopy was performed to monitor the increase in calcium levels in HUVEC after the addition of rPLP2. Selected frames of fluorescent images merged with DIC images with time elapsed between frames are shown. Scale bar indicates 5 μm.

### 2.3. rPLP2 leads to the exposure of phosphatidyl serine and propidium iodide positivity resulting in late apoptosis/necrosis

rPLP2 forms pore on cell membranes that could lead to lysis of cells. To analyze whether cell death was caused by lytic or non-lytic mechanisms (i.e., necrosis or apoptosis), cells incubated with rPLP2 was monitored for exposure of phosphatidyl serine and uptake of propidium iodide.

Cells positive for Annexin V alone indicate apoptotic as it only indicates the exposure of phosphatidylserine without cell permeabilization. Whereas, cells positive only for PI were considered necrotic indicating permeabilization. However, Annexin V/PI double-positive cells were considered late apoptotic/necrotic. Treatment of the cells with rPLP2 showed a clear Annexin and PI dual positivity, confirming cell death (Figure 4A). Moreover, mean intensity graph also showed a clear difference between the control and treated cells (Figure 4B). This data indicates that rPLP2 causes late apoptotic/necrotic of MDCK cells. Epsilon toxin was used as a control indicating its similar activity to rPLP2 (Figure 4A).

**Figure 4:**
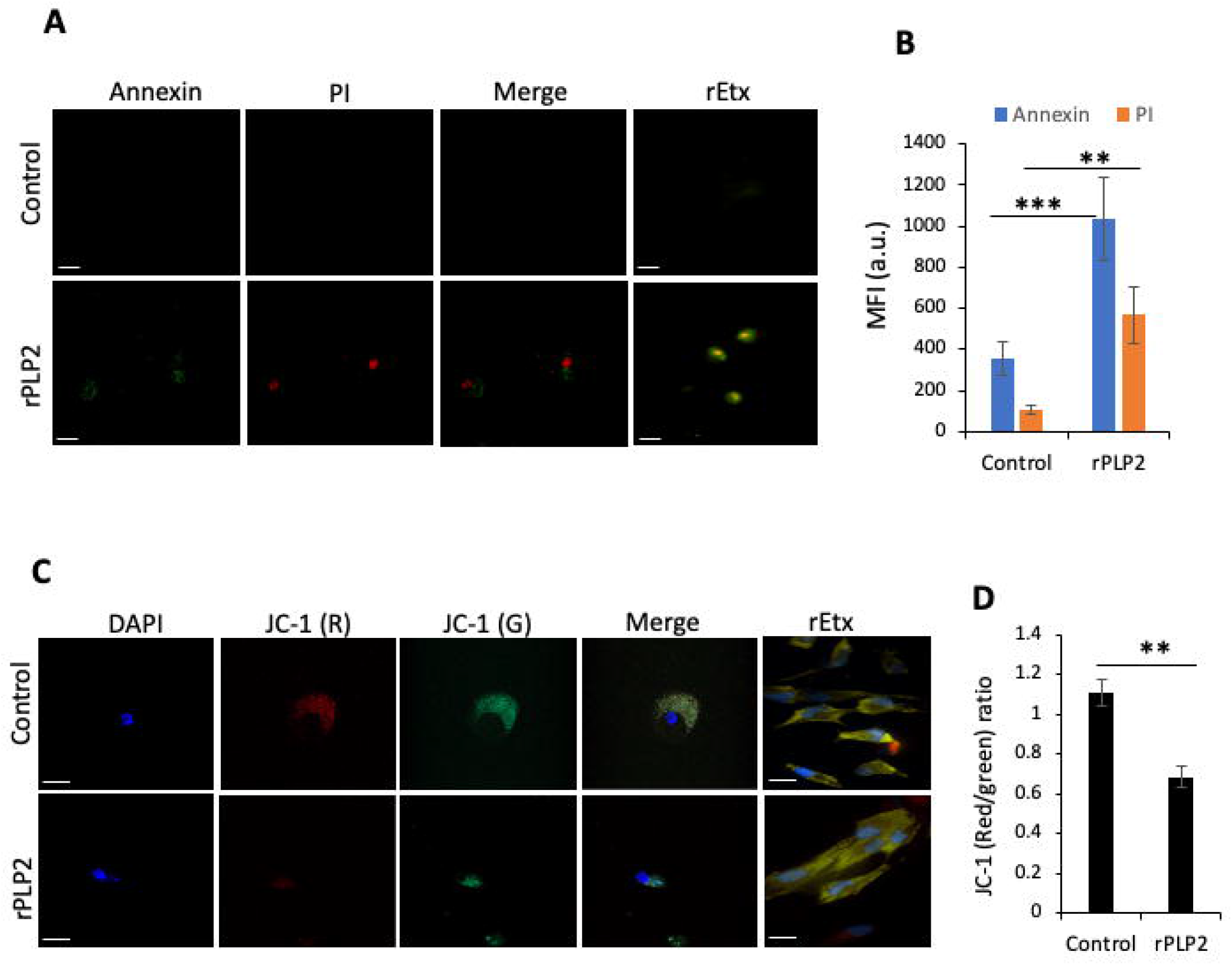
rPLP2 treated cells express markers of cell death. A) MDCK cells were treated in the presence or absence of rPLP2 and the Annexin V/PI staining was observed. Scale bar represents 20 μm. rEtx treated MDCK cells were taken as a control. B) The bar graph depicts the changes in the mean fluorescence for both Annexin and PI between control and treated groups. C) MDCK cells were loaded with JC-1 and the effect on mitochondrial membrane depolarization (ΨΔm) was observed. Scale bar represents 20 μm. rEtx treated MDCK cells were taken as a control. Error bars represents mean±SD. D) Bar graph depicts the red (aggregates)/green (monomers) ratio.

The accumulation of calcium by the mitochondria is a crucial step leading to the loss of mitochondrial membrane potential (ΨΔm) and energy collapse, in-turn causing cell death [14,15]. Calcium also contributes to the disruption of mitochondrial membrane potential, causing lower levels of ATP production [16]. Hence to monitor the effect of rPLP2 on mitochondrial disruption, we used JC-1 dye which is an indicator of ΨΔm. In the case of healthy cells, the mitochondria show a potential-dependent accumulation of JC-1 leading to the formation of J aggregates. This is followed by a shift in the emission from green (~529 nm) to red (~590 nm). However, in the case of depolarization, JC-1 exists in its monomeric form leading to green fluorescence. This depolarization is indicated by a decrease in the red/green fluorescence intensity ratio. Upon treatment of the cells with rPLP2, a rapid decrease in the mitochondrial membrane potential was observed as evident by the increase in the accumulation of green monomers (Figure 4C). A clear decrease in the red/green fluorescence intensity ratio could be observed as compared to the control cells which had intact potential (Figure 4D). Etx treated MDCK cells were used as a control.

### 2.4. rPLP2 increases the levels of intracellular reactive oxygen species in primary cells

Reactive oxygen species (ROS) are the highly reactive molecules produced within the cells. At low levels, they are considered to be important for the regulation of various processes including differentiation, proliferation, and cell death [17]. Excess production of ROS is lethal to the cells and can cause damage to DNA, proteins, lipids leading to the activation of cell death processes [18]. rPLP2 could enhance intracellular ROS generation that could lead to cell death. Hence, we monitored the ROS production in cells using a cell-permeant DCFDA dye (2’,7’-dichlorodihydrofluorescein diacetate). This dye, in the presence of ROS, exhibits higher levels of fluorescence, thus providing a reliable method for detection. In control cells, basal levels of ROS could be detected (Figure 5A). However, in rPLP2 treated cells higher levels of ROS could be observed. A bar graph shows a clear shift in the fluorescence intensity between the control and treated cells (Figure 5B). A similar increase in ROS could be detected in the MDCK cells exposed to Etx as described earlier [19].

**Figure 5:**
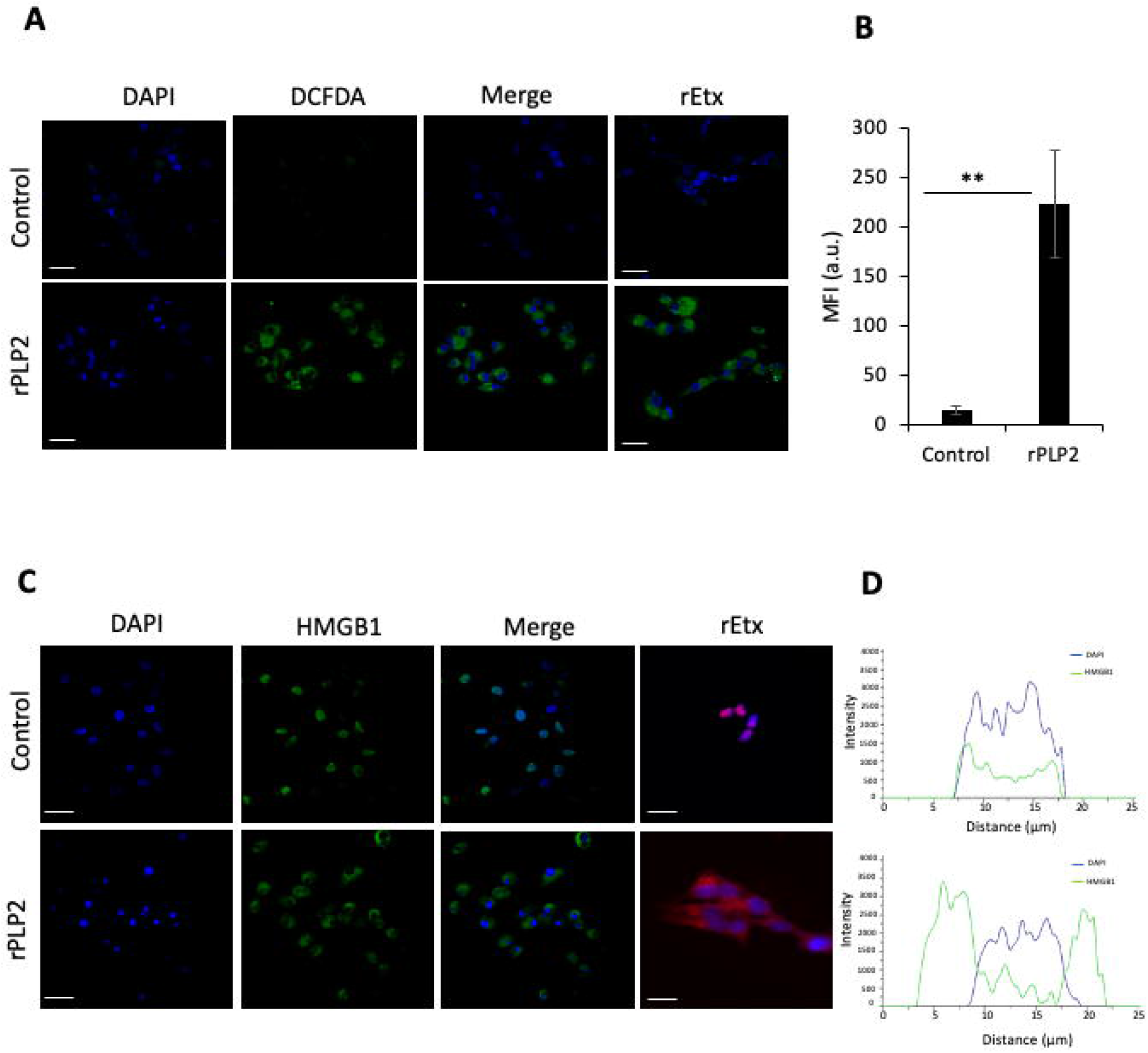
rPLP2 treated cells show ROS increase and HMGB1 translocation. A) Intracellular DCFDA levels were monitored in cells in the presence of absence of rPLP2. Scale bar indicated 20 μm. rEtx treated MDCK cells were taken as a control. B) Bar graph represents MFI for DCFDA between the control and treated groups. Error bars represents mean±SD. C) The localization of HMGB1 was monitored in the presence or absence of rPLP2. rEtx treated MDCK cells were taken as a control. D) Co-localization line profiles of DAPI (blue) and HMGB1 (green) are represented. Scale bars represents 20 μm.

### 2.5. rPLP2 treatment leads to the translocation of HMGB1 from the nucleus to the cytosol

Dying cells often display compromised membrane architecture, leading to the release of intracellular molecules that can act as endogenous adjuvants and foster inflammation [20]. HMGB1 is one such molecule and is integral to oxidative stress and downstream apoptosis or survival. In healthy cells, HMGB1 accumulates in the nucleus and is found at sites of oxidative DNA damage thus functioning in DNA repair [21]. However, in the case of necrosis, it functions as a cytokine mediator of inflammation. Studies indicate that ROS can release HMGB1 through calcium/calmodulin-dependent kinase-mediated signaling [22]. Hence, to check for the localization of HMGB1, we performed immunofluorescence assay in the presence or absence of rPLP2. In control cells, the HMGB1 was localized in the nucleus and no levels could be detected in the cytoplasm. Whereas upon treatment with rPLP2, HMGB1 migrated from the nucleus to the cytosol indicating cell necrosis (Figure 5C). Co-localization intensity plot analysis suggested the HMGB1 is present in the nucleus in control cells. Whereas, in the case of treatment, no co-localization could be observed indicating the shift from the nucleus to the cytoplasm (Figure 5D). Hence, both Etx and rPLP2 exhibit similar activity and lead to the necrosis of their target cells *in vitro*.

## 3. Discussion

The major functions of BBB are to mainly provide a controlled entry and exit of various molecules that infiltrate the brain through the EC tight junctions and its membrane proteins [23]. Through extensive studies in animal models, cultured cells, and patients, it has become evident that the dysfunction of BBB plays a critical role in the development of cerebral malaria. A number of factors, including inflammatory cytokines and ROS, have been implied in its disruption [24]. It is usually seen that in case of severe CM, the Pf-IRBCs adhere to the EC through specific interactions [5]. This could further lead to the blockage of cerebral vasculature in-turn blocking the normal blood supply and the movement of other molecules. Along with this, the toxins released by the Pf-IRBCs can also have a major role in the disruption of BBB. Overall, the parasite infection can cause cerebral edema along with haemorrhages and occlusions imposing deadly sequelae of neurological injury, promoting disease severity [25].

It is well known that *P. falciparum* secretes PLPs that play an important role in the progression of its life cycle. Our previous study [11] demonstrated that PLPs form pores on the erythrocytes leading to their lysis in the blood stage. Here in, we report that the rupture of Pf-IRBCs after schizogony leads to the release of PLPs, which in turn compromises Endothelial barrier integrity leading to the death of ECs.

To initially check the lethal effect of rPLP2 on mammalian cells, we performed MTT assay. As shown in the Figure1B, MDCK cells showed a dose-dependent lysis with an increase in rPLP2 treatment. Since the PFTs exposed cells often display changes in membrane architecture, we performed light microscopy. rPLP2 exposed cells showed changes in the cell morphology including cell rounding and membrane shrinkage (Figure 1C). Further, to observe the changes in real-time, we performed live-cell video microscopy. The rPLP2 treated cells initially displayed swelling at different sites on the membrane called blebs (Figure 2A and 2B). Further, these blebs appeared all over the cell membrane indicating features of apoptosis.

Calcium is a crucial player in the PFT induced toxicity and deregulation of its concentrations is critical for cell fate. PFTs cause a wide range of ion imbalances which involve Ca^2+^ influx and K^+^ efflux [26,27]. It is also observed that certain toxins mediate the release of Ca^2+^ from the ER [28,29]. Both the influx as well as the intracellular release lead to a drastic spike in Ca^2+^ levels. To ascertain this, we performed Fluo-4 AM staining to detect the changes in Ca^2+^ levels. Upon exposure to rPLP2, the levels of calcium rapidly increased within the cells which can be distinguished by the higher level of fluorescence in comparison to the control cells (Figure 3A and 3B).

Ca^2+^ influx is also associated with the dysfunction of mitochondria which occurs as a result of the calcium stimulation of the mitochondrial TCA cycle [16]. This can lead to ROS production, ATP depletion and resulting in death. Hence, to monitor the effect of rPLP2 on the cells, we used Annexin/PI dual staining. Our results indicate that rPLP2 mediates a late apoptotic/necrotic phenotype that can be deciphered from the Annexin/PI dual positivity (Figure 4A). Further downstream analysis indicated a drastic reduction for ΨΔm in cell treated rPLP2 compared to the control cells which had intact potential (Figure 4C and 4D).

ROS production is one of the markers for the overstimulation of the mitochondrial TCA cycle. They can further permeabilize the mitochondria by positive feedback [30] as well as disrupt the lysosomes [31]. Many studies have determined the role of ROS as a mediator for cell death in the case of PFTs [27,32]. Increase in levels of ROS also causes disruption in blood brain barrier during cerebral malaria. In this study, we could also clearly detect increased levels of ROS in the rPLP2 treated cells (Figure 5A). Importantly, malaria patients have been reported to have a lower catalase activity than healthy controls but a higher SOD activity, thus resulting in the accumulation of H_2_O_2_. Moreover, it is reported that free heme released from the hemoglobin of ruptured parasites infected red cells may continuously undergo autoxidation, producing super oxide, which dismutates into H_2_O_2_ and is a potential source for subsequent oxidative reactions in endothelial cells [33].

It is well known that increased ROS leads to DNA damage and decreased energy production [34]. All these factors finally lead to the death of the target cell. After the cellular membrane damage, the release of intercellular contents including HMGB1 is a cause for inflammation in necrosis [35]. The depletion of ATP occurs due to the overactivation of PARP causing necrosis. ROS leads to DNA damage leading to activation of PARP which further phosphorylates HMGB1 [36]. Upon staining for HMGB1, we found the levels were largely located in the nucleus in control cells (Figure 5C). However, a clear translocation from the nucleus to the cytosol (Figure 5C and 5D) was evident in the treated cells indicating necrosis.

Taken together, we provide substantial evidence that the rupture of Pf-IRBCs after schizogony leads to the release of PLPs, which in turn compromises BBB integrity. The addition of purified rPLPs forms pores on EC leading to an increase in intracellular Ca^2+^ and subsequent cell death. This study for the first-time sheds light on the role of PLPs on EC in the context of CM. Further, these PLPs can be attractive targets for designing inhibitors that can lead to enhanced cellular protection.

## 4. Materials and Methods

### 4.1. Cell culture

The Madin-Darby canine kidney (MDCK) cell line was maintained in DMEM (Gibco) supplemented with 10% FBS (Gibco) and 1% Penicillin/streptomycin (Gibco). Human umbilical vein endothelial cells (HUVEC) were cultured in EGM2 media (Lonza) supplemented with growth factors (Lonza).

### 4.2. Recombinant PLP2 and Etx purification

The purification of rPLP2 [11] and rEtx [37] was carried out as described previously. In general, the codon optimized gene encoding for rPMDs was subcloned into PET28a (+) and protein expression in *E.coli* cells was induced with 1 mM isopropyl-β-D-thiogalactoside (IPTG) (Sigma). The his-tagged proteins were purified using Ni-NTA chromatography and the concentration was determined using the BCA estimation kit (Pierce).

### 4.3. Cell viability

MDCK cells were cultured in 96-well plates at a density of 0.2×10^6^ cells/well to analyze the effects of purified rPLP2 or rEtx. Serial dilutions of the proteins in incomplete DMEM (iDMEM) or phosphate buffer were incubated with the cells in triplicates for 1 h at 37°C. The control wells had an equal volume of media. Cell viability was measured using MTT [3-(4,5-dimethylthiazol-2-yl), 2,5diphenyltetrazolium bromide] (Himedia) and the absorbance was read at A_550_ using Biorad iMark microplate reader (Biorad). Untreated cells were taken as 100% cell survival, and the viability for Etx-treated cells was calculated accordingly.

### 4.4. Live-cell video microscopy

MDCK or HUVEC grown on glass bottom culture dishes (ibidi) were treated with the rPLP2 and the morphological changes were observed in real-time using Nikon confocal microscope (Nikon A1R). For calcium influx, the cells were loaded with Fluo-4AM (Thermo Fischer Scientific) dye, followed by two washes to remove the excess dye. rPLP2 was added to the cells and the time-lapse images were captured using a 1.4 numerical aperture lens and analyzed by Nikon ES elements software.

### 4.5. Annexin-PI staining

MDCK cells were treated with the protein and incubated for 1 h at 37°C. The cells were washed with PBS and stained with Annexin V-FITC (Thermo Fischer Scientific) and PI (Thermo Fischer Scientific) and imaged. Similarly, the mitochondrial membrane potential indicator JC-1 (Thermo Fischer Scientific) staining was carried out after incubation with the protein for 1 h using the manufacturer protocol.

### 4.6. Mitochondrial membrane potential and ROS detection assays

The cells were grown on the glass bottom culture dishes (idbi) and incubated at 37°C overnight. Once the desired confluency was reached, the cells were incubated with the protein. After 1 h, the cells were washed and stained with JC-1 (Thermo Fischer Scientific) followed by imaging. Similarly, the DCFDA (2-7-dichlorodihydro fluorescein di-acetate) (Thermo Fischer Scientific) for ROS detection was carried out after incubation with the protein for 1 h using the manufacturer protocol.

### 4.7. HMGB1 assays

MDCK cells were grown on the sterile coverslips (Corning) and incubated for at 37°C overnight. The cells were treated with the protein for 1 h, and the coverslips were washed and fixed with 2.5% paraffin, followed by permeabilization. HMGB1 (Thermo Fischer Scientific) was used as a primary antibody followed by the addition of anti-rabbit Alexa 488 or anti-rabbit Alexa 594 (Thermo Fischer Scientific) as the secondary antibody. The coverslips were mounted on a glass slide using DAPI antifade (Thermo Fischer Scientific) and imaged under the confocal microscope.

### 4.8. Statistical analysis

Student’s t-test was performed where ever applicable. P-values of <0.05, <0.01 and <0.005 were considered significant (denoted by *, ** and ***) respectively. Results represent the mean ± SD of three independent experiments.

## Author contributions

A.S. performed the recombinant protein based *in vitro* assays. A.S (Akriti Srivastava), R.H. assisted in the protein and microscopy related work. S.G. performed the live cell imaging for blebbing and calcium uptake in HUVEC. A.S. and S.G. analyzed the data and wrote the manuscript. S.P. helped in the experimental analysis. L.G. provided the Epsilon toxin for the work. S.S. conceptualized and designed the entire study. All the authors read and approved the manuscript.

## Acknowledgments

A.S, A.S (Akriti Srivastava), R.H, are supported by the Shiv Nadar University fellowship. S.P is grateful for the support from the Shiv Nadar Foundation. The authors would like to thank AIRF for access to live-cell microscopy at JNU.

